# An Algorithmic Barrier to Neural Circuit Understanding

**DOI:** 10.1101/639724

**Authors:** Venkatakrishnan Ramaswamy

## Abstract

Neuroscience is witnessing extraordinary progress in experimental techniques, especially at the neural circuit level. These advances are largely aimed at enabling us to understand how neural circuit computations mechanistically *cause* behavior. Here, using techniques from Theoretical Computer Science, we examine how many experiments are needed to obtain such an empirical understanding. It is proved, mathematically, that establishing the most extensive notions of understanding *need* exponentially-many experiments in the number of neurons, in general, unless a widely-posited hypothesis about computation is false. Worse still, the feasible experimental regime is one where the number of experiments scales sub-linearly in the number of neurons, suggesting a fundamental impediment to such an understanding. Determining which notions of understanding are algorithmically tractable, thus, becomes an important new endeavor in Neuroscience.

---

Neuroscience is making remarkable ongoing progress in experimental techniques, at the present time. At the neuronal and network level, advances include the ability to both image activity in (*1, 2*), as well as activate/silence subsets (*3–5*) of neurons in-vivo, all-optically (*6*), in awake, behaving animals. Already, such techniques are being deployed to study (nearly) whole-brain neural activity at cellular resolution in C. elegans (*7*), hydra (*8*) & adult Drosophila (*9*) and to combine imaging of whole-brain neural activity with optogenetic perturbations of arbitrary subsets of neurons in larval zebrafish (*10*). Likewise, connectomes of the nematode C. elegans (*11*) and the larval tadpole C. intestinalis (*12*) have been fully reconstructed, as have significant portions of the larval fruitfly D. melanogaster (*13*).

Indeed, ongoing national initiatives in the United States (*14*), Japan (*15*), China (*16*) and Korea (*17*) have endeavored to accelerate the development of these (and other) neurotechnologies. The general premise has been that such technologies will ultimately enable us to reach the goal of understanding how networks of neurons mechanistically perform computations that lead to specific behaviors.

Understanding how the activity of neurons in circuits mechanistically causes behavior will require^1^ perturbational experiments at high spatial and temporal resolution that contemporary techniques are allowing us to approach. Efficient algorithms are needed that will seek to prescribe the smallest number of experiments necessary (that scale as a function of the number of neurons in the network) to establish such an understanding. An experiment, for example, might entail perturbing activity of a chosen subset of neurons, while concurrently imaging neural circuit activity and attempting to elicit behavior. The specifics of the next experiment prescribed by the algorithm could depend on the outcome of the current experiment. We are thus treating the problem of experiment design as an algorithmic problem^2^, invoking tools from Computer Science. Theoretical Computer Science has known, from over half a century of work, that some problems have fast algorithms whereas certain others provably require intractably many steps of computation to solve, in general. It is as yet unclear in which class of problems, those pertaining to empirically understanding neural circuits, fall in. Previous work (*18, 19*) has suggested that combinatorial explosion in number of interactions (among other considerations) might present a challenge to this end. Combinatorial explosion in the solution-space does not necessarily imply algorithmic intractability^3^, and therefore the question has remained largely open.

Here, using techniques from Computational Complexity Theory, we ask what is the smallest number of experiments *necessary*, in general, in order to arrive at a comprehensive empirical understanding of neural circuit computations that lead to a fixed behavior, in a hypothetical experimental setting. We find that no general algorithms exist to solve this class of problems, that always use sub-exponential number of experiments in the number of neurons, unless^4^ the complexity class 𝒫 = 𝒩𝒫 ^5^. This is the case even if the connectome is known. Performing exponentially-many experiments in the number of neurons would lead one to require more experiments than the estimated number of atoms in the observable universe, even for modest-sized nervous systems – rendering it an impracticable undertaking. To make matters worse, we demonstrate, using data and estimates, that the feasible regime of experiments – for most nervous systems of interest – is one where the practicable number of experiments that can be performed, scales *sub-linearly* in the number of neurons. This remarkable gulf between the worst-case and the feasible suggests a hitherto unappreciated, fundamental *algorithmic* barrier to understanding mechanistic computation in neural circuits. This algorithmic barrier is the proposition that, in spite of having adequate technological tools, we might lack tractable algorithms that can use those tools to distill aspects of empirical understanding of neural circuit computation. As a result, it becomes important to characterize the class of questions about mechanistic circuit computation that can be established empirically, in a tractable manner.

The experimental setting we consider here is one where we have an individual^6^ animal (that has been trained to be) doing a certain behavioral task and we wish to obtain a comprehensive mechanistic understanding of how its neural circuits (acutely) cause this behavior to be manifested at the present point in time^7^. This understanding must be a causal account and also encompass the degeneracy that has been associated with neural computations (*20*). We assume the existence of a behavioral readout that the experimenter is interested in^8^, which is a way of objectively partitioning possible behaviors into two classes, namely cases where the behavior is “correctly” expressed and cases where it isn’t. For instance, a possible task could be an odor recognition task, wherein the experimenter’s goal is to obtain an understanding of how mechanistic computation in neural circuits currently causes the animal to recognize the odor and perform the correct behavior to indicate the same. For the sake of analysis, we will assume that we have access to its entire nervous system, where we have the ability to, for example, image activity in and stimulate/silence arbitrary subsets of neurons and we know its connectome, although the main result is largely independent of the specific experimental capabilities/technology at hand.

What does it mean to understand mechanistic computation in neural circuits? There is, as yet, no standard definition of understanding in this context and conceivably, there exist multiple concomitant descriptions constituting notions of understanding that might, for example, include details spanning different spatial/temporal scales. We wish to have our theory be applicable to a wide variety of such notions. A characteristic of understanding is the ability to answer questions about the subject of understanding, in short order. Accordingly, the most extensive notions of understanding – which we will call *comprehensive notions of understanding* – ought to enable us to so answer certain basic questions about mechanistic computation in the circuit leading up to behavior. Here, we will propose *six* such questions that involve determining certain types of subsets of neurons that (causally) participate in computations leading to the said behavior. Furthermore, we show that no general tractable algorithms exist to establish notions of understanding that allow us to answer any of these questions in short order.

Next, we provide formal definitions that correspond to commonly-held notions of circuits being sufficient/necessary for the expression of a behavior, although these specific terms themselves have been deprecated (*21*). While the definitions are mostly posed in terms of silencing neurons, the circuit interrogation experiments themselves are free to stimulate or silence neurons (e.g. optogenetically).

The first definition corresponds to the notion of a sufficient circuit. Since there could concurrently exist multiple such sufficient circuits, each of them may be considered degenerate (*20*).

## Definition 1

(Degenerate Circuit). *A subset N of neurons is said to constitute a degenerate circuit for a behavior B, if B can be successfully elicited with the silencing of all neurons, except those in N*.

In the definition above, whether the behavior is successfully elicited or not is determined with respect to a fixed behavioral readout, as discussed earlier. Now, there always exists at least one degenerate circuit – the entire network – although, in practice, there likely exist many degenerate circuits. Presence of a neuron in a degenerate circuit does not necessarily imply a causal role for it^9^.

Degenerate circuits have been implicitly identified in Neuroscience for long via lesion studies and more recently via local cooling, pharmacological and optogenetic/chemogenetic/thermogenetic silencing, wherein the behavior in question persists in spite of silencing a number of neurons. Interesting recent cases have demonstrated the existence of degenerate circuits that exclude regions of the brain previously thought to be crucial for the studied behavior. For example, a non-dexterous motor task (*22*) does not require the motor cortex and likewise, a whisker-based object detection task (*23*), has been shown to not need the barrel cortex. We now define another notion.

## Definition 2

(Minimal Degenerate Circuit (MDC)). *A subset N of neurons is said to constitute a minimal degenerate circuit for a behavior B, if N constitutes a degenerate circuit for B and furthermore no proper subset of N is a degenerate circuit for B*.

Again, there exists at least one MDC with respect to each behavior, although there are likely many. In the context of a particular MDC, each neuron in it is necessary to elicit the behavior, since silencing it abolishes the behavior.

## Definition 3

(Minimum Degenerate Circuit). *A subset N of neurons is said to constitute a minimum degenerate circuit for a behavior B, if N constitutes a minimal degenerate circuit for B of the smallest size*.

The notion of a Minimum Degenerate Circuit is congruent with the goal of finding and characterizing the the most parsimonious circuit that is sufficient to cause the behavior.

Next, it is interesting to determine subsets of neurons, which when silenced are guaranteed to abolish the behavior (with respect to the said readout).

## Definition 4

(Vital Set). *A subset N of neurons is said to constitute a vital set of neurons for a behavior B, if N has a non-empty intersection with every degenerate set for B*.

It follows that if a vital set of neurons is silenced, then no degenerate set can be concurrently active and therefore the behavior cannot be correctly elicited, according to the readout.

## Definition 5

(*k*-Vital Set). *A vital set of size k is called a k-vital set*.

A 1-Vital set would in fact correspond to the intersection of all degenerate sets.

As with degenerate circuits, we define a notion of minimality for vital sets that corresponds to the commonly-held notion of a set of neurons being necessary for expression of the behavior.

## Definition 6

(Minimal Vital Set). *A subset N of neurons is said to constitute a minimal vital set of neurons for a behavior B, if N is a vital set and no proper subset of it is a vital set*.

As with degenerate sets, putative vital sets have been identified in cases, where silencing a set of neurons abolishes the behavior in question. Remarkably, there also exist examples of putative minimal vital sets being identified experimentally (*24–26*).

Following the definitions, we now state six questions about mechanistic circuit computation leading to behavior. We argue that notions of understanding that cannot enable us to answer any of these questions, in short order, cannot claim to be comprehensive notions of understanding. Subsequently, we show that none of these questions can be answered with sub-exponential number of experiments in the number of neurons, in general, unless 𝒫 = 𝒩𝒫. The first three questions pertain to subsets of neurons *sufficient* to express behavior and the last three questions are about subsets of neurons that are *necessary* to cause behavior. The questions^10^ are:

1. Determine a Degenerate Circuit of size *k* or indicate if none exists of that size.
2. Determine a Minimal Degenerate Circuit of size *k* or indicate if none exists of that size.
3. Determine a Minimum Degenerate Circuit.
4. Determine the set of 1-Vital sets (or indicate if there are no 1-Vital sets).
5. How many minimal *k*-Vital Sets exist?
6. Determine a minimal *k*-Vital Set, or indicate if none exists of that size.

More precisely, we define a notion of understanding to be comprehensive, if from its description, one can quickly (i.e. in number of steps that scale as a polynomial in the number of neurons) answer any of the above questions. The definition, and the methods that follow, utilize the notion of a *reduction*^11^, which is fundamental in Theoretical Computer Science.

For each of the six problems, we establish a reduction from a known 𝒩𝒫-complete problem (Supplementary Text). This would entail providing a “recipe” that takes any instance of said 𝒩𝒫-complete problem and quickly constructs a neural circuit with the guarantee that if the said question about the neural circuit can always be answered efficiently, the said 𝒩𝒫-complete problem can be solved likewise. This implies that no algorithms exist for these questions that always take sub-exponential number of experiments, unless 𝒫 = 𝒩𝒫. In a sense, this is a way of raising the stakes – implying that a general efficient algorithm for the said neural circuit question would lead to general efficient algorithms for hundreds of computational problems, for which no such algorithms have been known to exist yet.

An important caveat is that this analysis – as is typical in Computational Complexity Theory– is of the worst-case. That is, the theory implies that there is a sub-class of neural circuits that provably require exponentially-many experiments for the said problems, unless 𝒫 = 𝒩𝒫. Worryingly though, this sub-class of circuits is a rather simple-looking one, that maps stimulus to response, while potentially being mediated by a smaller degenerate circuit. It is natural, therefore, to expect that circuits performing seemingly more complex computations might be at least as hard to understand comprehensively. On the other hand, even for circuits that may lie outside this sub-class and use sub-exponential number of experiments for the said problems, in order to be realizable, such algorithms, in fact, can use only sub-linear number of experiments, as we argue next – an onerous requirement indeed. This extraordinary gap between the worst-case and the realizable is at the heart of this algorithmic barrier to establishing a comprehensive empirical understanding of neural circuits.

While, performing exponentially-many experiments in the number of neurons is clearly infeasible, what is the feasible regime of experiments? To obtain a rough upper bound, we consider how many experiments might one hope to perform during the lifetime of an individual animal and how this compares to the number of neurons in it. Using estimates of average lifespans and number of neurons, we demonstrate (Figure 2A) that for most organisms of interest, we can only perform sub-linear^12^ number of experiments in the number of neurons. Even if we had neural proxies for behavior, which manifest in – e.g. a 100ms – the feasible experimental regime is not dramatically different (Figure 2B). A sub-linear experimental regime is a particularly stringent prospect, even for notions of understanding that may not be comprehensive. Especially notable is the extraordinary gap between the worst-case (exponential) and feasible (sub-linear).

**Figure 1:**
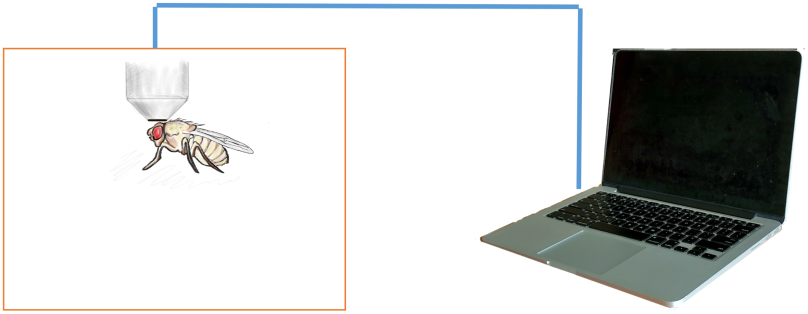
A schematic visualizing an example of an experimental setting that is considered here. Given an individual organism performing a certain behavior, we wish to comprehensively understand how its neural circuits mechanistically cause the behavior to be manifested. Specifically, the question is how many neural circuit interrogation experiments might be necessary to establish such an understanding.

**Figure 2:**
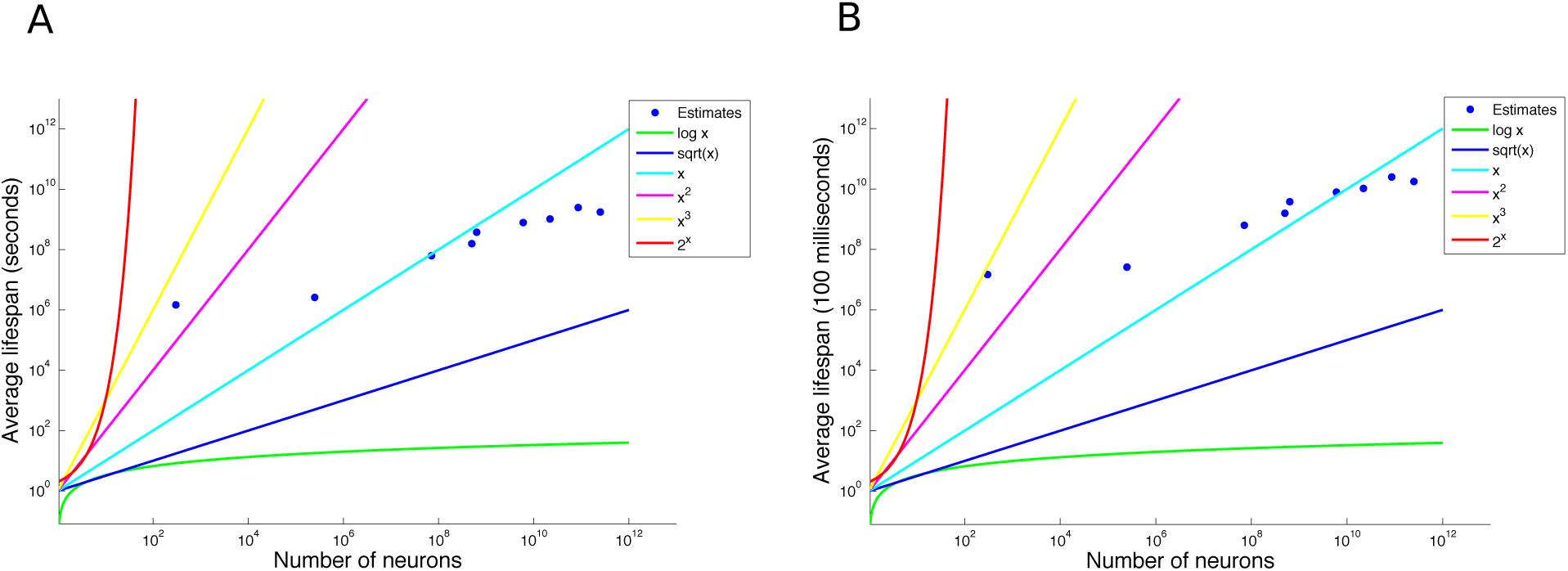
(A) A log-log plot of estimates of the average lifespan as a function of (estimated) number of neurons in the (central) nervous system for C. elegans, fruit fly, mouse, octopus, marmoset, Chimpanzee, Rhesus monkey, Human, and Elephant. Superimposed, is the time it would take to finish an experimental protocol that is logarithmic, linear, quadratic, cubic and exponential in the number of neurons, assuming each experiment takes a second (corresponding to the behavioral timescale) and that there is no gap between successive experiments – both conservative assumptions. The plot suggests that running even a linear number of such experiments for most organisms would exceed their expected lifespan. (B) The corresponding plot, assuming each experiment takes only 100 ms. The feasible regime does not dramatically improve.

To close, empirically understanding how neural circuits mechanistically cause behavior appears to be a pre-requisite, for distilling the principles that govern their operation, or to validate said principles post hoc. The results here suggest that in addition to current technological impediments, there exists another fundamental barrier – an *algorithmic* one – to neural circuit understanding. Specifically, going forward, causal understanding of neural circuits will likely be frequently driven by algorithmic and theoretical constraints, rather than limitations in experimental methods.

Consequently, it becomes important to determine which notions of understanding are algorithmically feasible to establish in what contexts. This can be done in two related, but distinct ways. On the one hand, we need to ask what notions of understanding have algorithms that are guaranteed to take a small number of experiments to establish, no matter what class of circuits are involved. For example, in the Supplementary Text, we define the (non-comprehensive) notion of a Quasi-minimal degenerate circuit and describe an algorithm to determine an instance of it, which always needs only a logarithmic number of experiments in the number of neurons. For a brain with a hundred billion neurons, one would need approximately 40 experiments with this algorithm – a tractable number. The second approach, is to tailor questions to the specific subclasses of neural circuits under study, wherein one is guaranteed to answer the said questions with a small number of experiments for the specific circuits under consideration, even if the question may not be so answerable, in general. This second direction will involve considerable theory and appears to be the more arduous, if fruitful route.

## Acknowledgments

The author wishes to thank Moshe Abeles, Kambadur Ananthamurthy, Arunava Banerjee, Upi Bhalla, Vishaka Datta, Konrad Kording, Arvind Kumar, Sahil Moza, Dinesh Natesan, Sriram Narayanan, Christos Papadimitriou, Sanjay Sane, Shuchita Soman and K. VijayRaghavan for useful discussions, Dhananjay Chaturvedi for assistance with a figure and Erik Peterson for comments on a draft of the manuscript.

## Funding

The work was supported by the Simons Foundation.

## Authors contributions

VR conceived the idea, performed the research & wrote the paper.

## Competing interests

The author declares that he has no competing interests.

## Data and materials availability

All data is available in the manuscript or the supplementary materials.

## Supplementary Text

### Preface

Here, we present material supporting the main text of said paper. Specifically, we include a more rigorous mathematical account of the methods and present a notion of a specific type of degenerate circuit along with an algorithm that uses only a logarithmic number of experiments in the number of neurons to determine it. We close with concluding remarks.

The exposition is mostly mathematically self-contained, if somewhat brisk; reading the main text first will ease the reading here. Some parts might require a familiarity with basic notions in Theoretical Computer Science in order to grasp completely, although we make an effort to provide intuition and pointers to expository literature.

### 1 Combinatorial explosion of solution-space versus algorithmic hardness

Previous work (*27, 28*) has suggested that combinatorial explosion in number of interactions (among other considerations) might present a challenge to understanding neural circuits. Combinatorial explosion in the solution-space does not necessarily imply algorithmic intractability.

We will now expand on this distinction between combinatorial explosion in the solution-space of a problem versus the question of existence of an efficient algorithm for determining the solution of the same problem. Specifically, the mere existence of the former does not imply that there is no efficient algorithm to solve the said problem. Several examples abound in Computer Science, where efficient algorithms used every day, solve problems with exponentially-large solution spaces. For instance, given a labeled graph, which could represent road networks where nodes represent locations and edges represent roads labeled with distances, the problem of finding the shortest path between any two nodes can be solved (using Dijkstra’s algorithm) in number of steps that scales as the square of the number of nodes. This is in spite of the fact that if there were a road between every pair of locations, there would be exponentially-many paths – any of which might a priori be the shortest – given a start node and a destination node. Indeed, variants of this algorithm find application every day in portable electronic map systems. As a result, demonstrating algorithmic hardness tends to be more difficult than showing that the solution spaces are dauntingly large. Conversely, a small solution space immediately implies an efficient algorithm, so showing a potentially large search space is a useful prelude to asking the algorithmic hardness question. In this work, our focus, specifically, is on the algorithmic question.

### 2 Definitions, reductions and proofs

#### 2.1 Counting experiments

On the face of it, it may seem that the issue of counting the number of such experiments is a difficult problem, since it might appear to intimately depend on the details of experimental capabilities at hand. Fortunately, Theoretical Computer Science has already confronted and largely solved this question decades ago, in the case of computational devices, where the number of steps might depend on hardware details. The solution, briefly, is to not count the precise number of steps, but to determine how the number of steps grows asymptotically with the size of the input. Roughly speaking, this corresponds to asking how the number of steps used/required scales as a function of the input size. Foundational theorems (*29*) in Computer Science show that, in effect, details of hardware implementations^1^, essentially manifest as constant multiplicative/additive factors in such an account. Therefore, the notion of computational problems *needing* a certain number of steps, in this asymptotic sense, is a well-defined and fundamental one and indeed a number of such specific results are now known for many problems, from over half a century of Theoretical Computer Science. For example, in order to sort a list of *n* elements, using only comparisons, there is a mathematical proof that one *needs* of the order of *n* log *n* number of steps, in general. The reader is referred to Chapter 1 of (*30*) for a more detailed exposition of this issue.

#### 2.2 Problem instance

We formalize the problem instance to roughly correspond to experimental capabilities currently present (or being developed). That said, the result is largely independent of the specific experimental capabilities at hand, as we will elaborate. Indeed, the reductions would go through even if we had a precise simulation of the entire neural circuit that we could manipulate, at will.

For our purposes, next, we define a neural circuit, formally, as one, whose connectome is known^2^ and whose neurons we could manipulate (e.g. optogenetically), while measuring their activity (e.g. by 2-photon Calcium imaging or using a voltage indicator) and attempting to elicit a certain behavior with an associated binary behavioral readout.

##### Definition 1

(Neural Circuit). *A Neural Circuit 𝒩 is a 2-tuple 〈G, E : r × S_e_ × S_i_ → A × B_r_〉*

*where*

1. *G(V_G_, E_G_) is the connectome, as represented by a directed graph, where V_G_ is the set of neurons and E_G_ is the set of directed edges between them*.
2. *E(·, ·, ·) is the “experiment” function which given*

a. *r: an encoding of the state*.
b. *S_e_: a spatiotemporal matrix specifying how each neuron is (e.g. optogenetically) stimulated during the course of the experiment.*
c. *S_i_: a spatiotemporal matrix specifying how each neuron is (e.g. optogenetically) inhibited during the course of the experiment.* *gives us back*

a. *A: a spatiotemporal matrix that contains the activity of all the neurons during the course of the experiment*.
b. *B_r_* = {0, 1}*: the binary behavioral readout*.

According to the above definition, an experiment would correspond to the act of invoking the function *E*(*·*) once. An algorithm, given the connectome *G* and access to this function, might be tasked with answering specific questions about computation in this circuit. To this end, it has the ability to demand arbitrary experiments, with the possibility of subsequent experiment(s) adaptively depending on the outcome of the current experiment. We will end up showing that, in spite of such strong experimental capabilities and the ability to use them at will, there are questions which require exponentially-many experiments to answer^3^, in general, no matter how “clever” the algorithm.

The definition itself is set up to reflect, for example, a contemporary all-optical experiment on an awake behaving animal. However, this specific experimental setup isn’t particularly crucial for the results to hold, as we now elaborate.

Algorithms, effectively, are steps prescribed to solve a specific problem. A sequence of experiments, as exemplified above, also implicitly specifies an algorithm. That’s because each experiment above can be translated into a sequence of steps (with the number of steps, in this case, being linear in the number of neurons). We show here a number of questions about neural circuits that require exponentially-many steps in the number of neurons^4^, in general. Now, it cannot be, that answers to those questions can always be determined with sub-exponential number of experiments, for otherwise, such a schedule of experiments could be used^5^ to construct an algorithm that could always use only a sub-exponential number of steps as well, resulting in a logical contradiction. Indeed, any experimental setup – present or future – where an experiment can be translated into a polynomial number of steps in the number of neurons, would be one where these results hold. Also, it follows that the results would hold in a more restrictive experimental set-up than above. An example would be an experimental set-up where one could only perturb a small subset of neurons that might be in the field-of-view of the microscope in the current experiment. In short, the results have significantly broad applicability, including the case where we may have an exact simulation of the circuit that we could manipulate at will.

Notions of degenerate circuit, minimal and minimum degenerate circuit and the corresponding problem have been defined in the main text, which we repeat here, for completeness. The remainder of the section is a formalization of the material already covered in the main text. For this reason, the pace of exposition will be somewhat brisk.

##### Definition 2

(Degenerate Circuit). *A subset N of neurons is said to constitute a degenerate circuit with respect to the present behavioral readout, if the readout can be successfully elicited with the silencing of all neurons, except those in N*.

##### Definition 3

(Minimal Degenerate Circuit (MDC)). *A subset N of neurons is said to constitute a minimal degenerate circuit with respect to the present behavioral readout, if N constitutes a degenerate circuit with respect to the present behavioral readout and furthermore no proper subset of N is a degenerate circuit with respect to the present behavioral readout*.

##### Definition 4

(Minimum Degenerate Circuit). *A subset N of neurons is said to constitute a minimum degenerate circuit with respect to the present behavioral readout, if N constitutes a minimal degenerate circuit of the smallest size with respect to the present behavioral readout*.

##### Definition 5

(Vital Set). *A subset N of neurons is said to constitute a vital set of neurons with respect to the present behavioral readout, if N has a non-empty intersection with every degenerate set with respect to the present behavioral readout*.

It follows that if a vital set^6^ of neurons is silenced, then no degenerate set can be concurrently active and therefore the behavior cannot be correctly elicited, according to the readout.

##### Definition 6

(*k*-Vital Set). *A vital set of size k is called a k-vital set*.

##### Definition 7

(Minimal Vital Set). *A subset N of neurons is said to constitute a minimal vital set of neurons with respect to the present behavioral readout, if N is a vital set and no proper subset of it is a vital set*.

Before proceeding further^7^, we briefly review the notion of a reduction, which is a fundamental one in Theoretical Computer Science. Informally, a reduction is a recipe to quickly convert any instance of one problem (Problem A) into an instance of another problem (Problem B), such that a solution to Problem B can be quickly mapped back to a solution of Problem A as well. Therefore, the existence of an efficient algorithm for Problem B immediately implies the existence of an efficient algorithm for Problem A, via the reduction. More profoundly, if there exists no efficient algorithm for Problem A, the reduction implies that Problem B cannot have an efficient algorithm either; otherwise there would be a logical contradiction. In the description above, the words *quick* and *efficient* are, conventionally, synonyms for algorithms that take number of steps that scales roughly as some polynomial in the input size (which, for our purposes, will be proportional to the number of neurons). This is the usual convention in Theoretical Computer Science, for reasons that we will not get into here, and it is generally accepted that algorithms that need more steps than a polynomial function of input size are not tractable, except when the input size is very small. There exist multiple types of reductions. The ones we will be using in the definitions that immediately follow are called *Cook reductions*^8^, which assume the availability of a polynomial-time algorithm for Problem A that uses polynomially-many “function calls” to an algorithm for Problem B (such that the result of solving Problem B may be used to solve Problem A). This is a more general and liberal notion of a reduction unlike the *Karp reduction*^9^, which is for *decision problems*. A decision problem is a problem that takes in some input and always returns a Yes/No answer. Polynomial-time reductions between decision problems, wherein, a Yes answer for Problem B is mapped back to a Yes answer for Problem A and likewise a No answer for Problem B is mapped back to a No answer for Problem A is called a Karp reduction. We will use Karp reductions to prove that problems are NP-hard, by reducing a known NP-complete problem to the problem in question. The theory of 𝒩𝒫-completeness has historically been formulated for decision problems and therefore reductions that show 𝒩𝒫-completeness or 𝒩𝒫-hardness use Karp reductions. The six problems we list also involve what are called *search problems*, which is to say problems which involve searching for a specific type of object (e.g. “Determine a Minimal Degenerate Circuit of size *k* or indicate if none exists of that size.”). While search problems, technically, cannot be 𝒩𝒫-hard^10^, one could, in principle, reduce an 𝒩𝒫-hard problem to the search problem in question via a Cook reduction (See (*30*), Section 2.5, for more on this issue). This would imply that no sub-exponential algorithms exist for the said search problem, unless 𝒫 = 𝒩𝒫. Indeed, in many cases, we will first pose a decision version of the search problem and establish a Cook reduction from the decision version to the search problem, before proving that the decision version is 𝒩𝒫-hard.

We now expand and formalize a definition of neural circuit understanding, and then define comprehensive notions of understanding.

In order to be as general as possible (and this may seem abstract), we consider any problem that takes a neural circuit (as defined above) and produces an output (encoded as a string, traditionally) to constitute a notion of neural circuit understanding. In a sense, we would like to think of a notion of understanding as a certain form of a condensed distillation of data, from experiments performed on a single individual, with respect to a single behavioral readout. One might then consider using this distilled data to answer questions about mechanistic circuit computation in that individual, with respect to that behavioral readout. The obvious question, is what forms of distilled data should we seek to obtain and what questions will it enable us to answer. Secondly, of the questions it can enable us to answer, what questions can be answered in a timely manner?

The notion of understanding might simply be an answer to a certain question about the neural circuit or computation in it or something more complicated such as raw data from a small set of experiments. An example question would be determining the set of neurons with firing rate^11^ above a certain threshold. A notion of understanding that stores action potentials of all the neurons over the course of the experiment is one that can enable us to answer this question.

##### Definition 8

(Neural Circuit Understanding). *Any problem that takes as input a neural circuit and produces as output a string which is of polynomial length in the size of input is said to constitute a notion of Neural Circuit Understanding*.

Next, we distinguish certain types of neural circuit understanding as *comprehensive* notions of neural circuit understanding. The idea is to assert that if a notion of understanding claims to be comprehensive, then it ought to enable us to answer certain basic questions about mechanistic circuit computation with respect to the said behavioral readout. The intent here is not to come up with such an exhaustive list of questions, but only to list a few examples of such questions. We will go on to show that none of these questions have general sub-exponential algorithms, unless 𝒫 = 𝒩𝒫. As a result, no comprehensive notion of understanding, as defined below, can itself be established with sub-exponential number of experiments in general, unless 𝒫 = 𝒩𝒫.

##### Definition 9

(Comprehensive Neural Circuit Understanding). *A problem is said to constitute a notion of Comprehensive Neural Circuit Understanding if it constitutes a notion of neural circuit understanding and if there exists a Cook reduction from the one of the problems below to it*.

1. *Determine a Degenerate Circuit of size k or indicate if none exists of that size*.
2. *Determine a Minimal Degenerate Circuit of size k or indicate if none exists of that size*.
3. *Determine a Minimum Degenerate Circuit*.
4. *Determine the set of 1-Vital sets (or indicate if there are no 1-Vital sets)*.
5. *How many minimal k-Vital Sets exist?*
6. *Determine a minimal k-Vital Set, or indicate if none exists of that size*.

A remark about the formulation of the six problems, above, is in order. The problems above have been deliberately formulated in a way that ensures that the size of the solution itself is not exponential in the size of the input, for otherwise just stating the output would need exponentially-many steps in the worst case.

Next, we show that each of these six problems does not have algorithms that use sub-exponential number of experiments, in general, unless 𝒫 = 𝒩𝒫. To this end, the general strategy, often, will be to formulate the decision version of the stated problem, and establish a Cook reduction from that decision problem to the stated problem. The next step will then be to establish a Karp reduction from a known 𝒩𝒫-complete problem to this decision problem.

#### 2.3 Problem 1: Determine a Degenerate Circuit of size ***k*** or indicate if none exists of that size

We first formally state this question and it’s decision version.

##### Definition 10

(DC problem). *For a given positive integer k, determine a Degenerate Circuit of size k or indicate if none exists of that size*.

##### Definition 11

(DC (decision) problem). *For a given positive integer k, does there exist a Degenerate Circuit of size k?*

We now state the Cook reduction from the decision problem to the DC problem.

##### Proposition 1.

*DC (decision)* 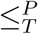 *DC*

The reduction simply entails solving the DC problem for the same inputs and examining if the solution returned a degenerate circuit of size *k* or if it is indicated that there is no such degenerate circuit; in the former case the decision version returns a *Yes* and in the latter case it returns a *No*.

Next, we prove that the DC (decision) problem is 𝒩𝒫-complete by establishing a Karp reduction from CLIQUE – a known 𝒩𝒫-complete problem – to DC (decision) and, furthermore, by showing that DC (decision) is in the complexity class 𝒩𝒫^12^. This implies that no general sub-exponential algorithms exist for any of the problems participating in the aforementioned reductions, unless 𝒫 = 𝒩𝒫.

The *clique* of an undirected graph is a subgraph of it, such that every pair of vertices in that subgraph has an edge between them. A clique that has *k* vertices is called a *k*-clique. The CLIQUE problem, which is a decision problem, seeks to determine if, for a given positive integer *k*, a given graph has a *k*-clique. CLIQUE has been shown to be 𝒩𝒫-complete. Establishing the reduction entails providing a “recipe” for turning *any* undirected graph into a neural circuit with the guarantee that the graph has a *k*-clique if and only if the constructed neural circuit has a degenerate circuit of a certain size. The proof below only describes the simplest construction. Afterward, we describe ways to further generalize the construction.

##### Lemma 1.

*DC(decision) is 𝒩𝒫-complete*.

**Figure S1:**
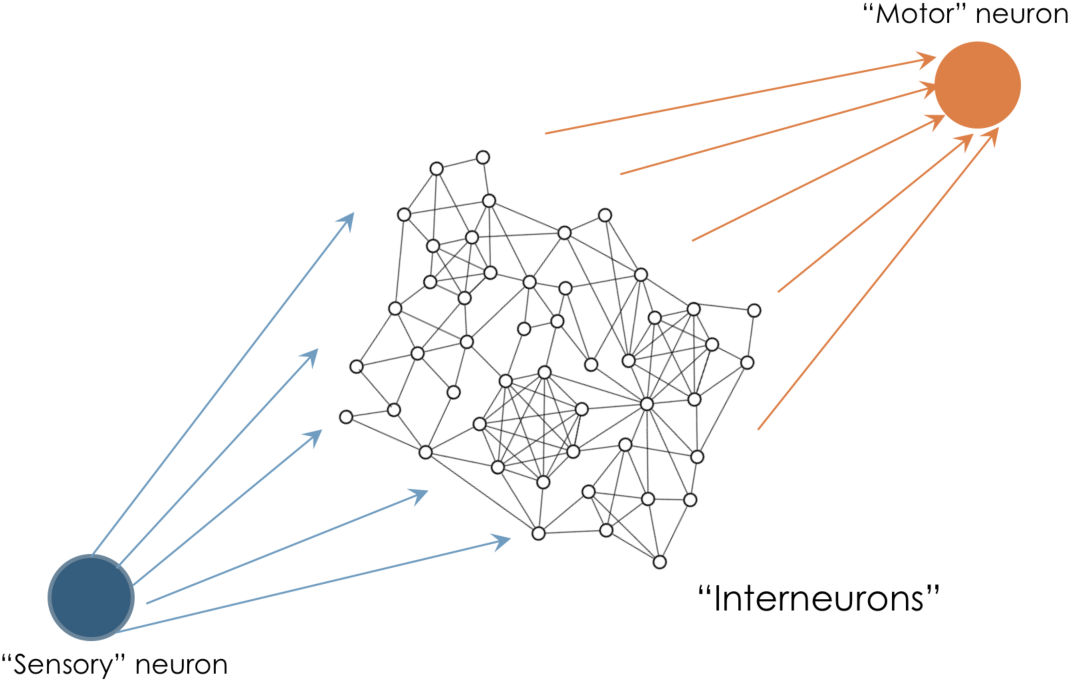
A schematic illustrating the neural circuit created by the reduction. The circuit has a sensory neuron, a motor neuron and an interneuron circuit which is constructed from the given undirected graph with the guarantee that this neural circuit has a (*k* + 2)-sized Degenerate Circuit if and only if the undirected graph has a *k*-clique.

*Proof.* We first detail the construction of the neural circuit given an undirected graph and a positive integer *k*, such that the neural circuit has a (*k* + 2)-sized DC if and only if the graph has a *k*-clique. The neural circuit has a single “sensory” and a single “motor” neuron. The sensory neuron signals the arrival of the pertinent stimulus by firing a single spike and the motor neuron, likewise, signals execution of the said behavior by firing a single spike (Figure S1). The “interneuron” circuit connecting the sensory and motor neuron is constructed from the undirected graph by having an excitatory (inter) neuron in place of each graph vertex and bi-directional connections^13^ between corresponding neurons whenever there is an edge between vertices. Every interneuron receives input from the sensory neuron and is in turn connected to the motor neuron. Conduction delays are the same within each pair of the following neuron types: sensory-inter, inter-inter and inter-motor. We consider here the state of the network, when all neurons start from quiescence. The synapses connecting the sensory neuron and the interneuron are strong, so that a single sensory neuron spike causes an interneuron spike. Interneuron-to-motor neuron synapses are somewhat weak; concurrent spikes from all interneurons is sufficient to cause the motor neuron to spike. However, if there is a pair of volleys^14^ of spikes from a set of *k* interneurons, the motor neuron (just about) spikes. Also, spiking from a set of less than *k* interneurons alone can, in no event, cause the motor neuron to spike. Finally, if an interneuron is not concurrently receiving input from the sensory neuron, it needs concurrent spiking from at least (*k* - 1) interneurons that are connected to it, in order to produce a spike.

As a result of the above construction, the neural circuit, ends up having the following properties: (a) The entire circuit forms a degenerate circuit for the behavior, with respect to the readout. (b) The undirected graph has a *k*-clique, if and only if the circuit formed by the corresponding neurons plus the sensory and motor neuron form a degenerate circuit of size (*k* + 2) for the behavior. (c) The circuit, by design, does not have a degenerate circuit of size less than (*k* + 2).

(a) follows immediately from the fact that concurrent spiking from all interneurons elicits a spike from the motor neuron; a spike from the sensory neuron is sufficient to elicit a spike from each interneuron. For (b), first consider the case when the graph has a *k*-clique. Consider the neural circuit, while silencing all interneurons except the ones corresponding to the vertices present in the *k*-clique. This circuit will lead to the motor neuron spiking when there is a sensory neuron spike. This is because the *k* “clique” neurons will fire a pair of volley of spikes, which are received by the motor neuron. The first volley comes about because of the sensory neuron spike. The second volley is a result of each neuron receiving a spike from *k* - 1 other “clique” interneurons, due to which it fires a spike. Now, consider the converse, namely the case where the graph has no *k*-clique. Now, every degenerate circuit has to have both the sensory and motor neuron present, since the circuit starts from the neurons being in the quiescent state. Now consider a circuit obtained by silencing all interneurons except *k* of the interneurons. From the construction, this circuit can elicit a motor neuron spike, if and only if *k* interneurons can send a pair of volley of spikes. However, the second volley cannot have *k* spikes. This is because there is at least one neuron among the *k* non-silenced interneurons that is receiving input from less than *k* - 1 other non-silenced interneurons^15^. As a result, that neuron cannot participate in the second volley, which ends up comprising less than *k* spikes, which prevents the motor neuron from spiking, in turn. (c) follows from the construction of the neural circuit; specifically, spiking from less than *k* interneurons is insufficient to cause the motor neuron to spike.

The above reduction implies that DC(decision) is 𝒩𝒫-hard. The problem also is in the complexity class 𝒩𝒫. The certificate, in this case is the degenerate circuit itself and one can check in linear time (using 1 experiment), whether the claimed subset of neurons indeed forms a degenerate circuit. As a result, DC(decision) is in the complexity class 𝒩𝒫 and this implies that it is 𝒩𝒫-complete.

Also, observe that the size (*k* + 2) degenerate circuit generated in the reduction, is both a minimal DC as well as a minimum DC. This implies that the same reduction establishes that the decision version of those corresponding problems also become 𝒩𝒫-hard, as we will elaborate in the next two subsections.

We now briefly remark on ways to further generalize the construction. For example, the construction has bi-directional connections between every pair of interneurons that is connected, which is not thought to be the case, physiologically. Likewise, all interneurons in the above construction are excitatory. Both these can be mitigated by the following straightforward addition to the above construction. Introduce arbitrary number of auxiliary interneurons (excitatory/inhibitory) with the constraint that each of them may receive connections or send out connections to at most (*k* - 2) other interneurons, which may be chosen arbitrarily. The inhibitory neurons do not send connections to the motor neuron. As a result, one has a neural circuit where the proportion of connections which are bidirectional in relation to unidirectional connections can be made arbitrarily small. Similarly, the circuit could also have proportion of excitatory and inhibitory neurons consistent with current estimates. The constraint described above on the number of connections ensures that the circuit has a (*k* + 2) sized degenerate circuit, if and only if the undirected graph has a *k*-clique.

#### 2.4 Problem 2: Determine a Minimal Degenerate Circuit of size ***k*** or indicate if none exists of that size

We first formally state this question and it’s decision version.

##### Definition 12

(MinimalDC problem). *For a given positive integer k, determine a Minimal Degenerate Circuit of size k or indicate if none exists of that size*.

##### Definition 13

(MinimalDC (decision) problem). *For a given k, does there exist a Minimal Degenerate Circuit of size k?*

We now state the Cook reduction from the decision problem to the DC problem.

##### Proposition 2.

*MinimalDC (decision)* 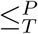 *MinimalDC*

As before, the reduction entails solving the MinimalDC problem and examining if the solution returned a circuit of size *k* or if it is indicated that there is no such degenerate circuit; in the former case the decision version returns a *Yes* and in the latter case it returns a *No*.

Next, we prove that the MinimalDC (decision) problem is 𝒩𝒫-hard by establishing a Karp reduction from CLIQUE. This implies that no general sub-exponential algorithms exist for any of the problems participating in the aforementioned reductions, unless 𝒫 = 𝒩𝒫.

##### Lemma 2.

*MinimalDC(decision) is 𝒩𝒫-hard*.

The reduction that was used to establish that DC(decision) is 𝒩𝒫-hard also turns out to imply that MinimalDC(decision) is 𝒩𝒫-hard. This is because, as proved in that construction, the (*k* + 2) sized degenerate circuit is also a minimal one, since the constructed circuit can, under no circumstances, have a (*k* + 1)-sized or smaller degenerate circuit.

Note that we are not asserting that MinimalDC(decision) is 𝒩𝒫-complete, since it is unclear if this problem is in the complexity class 𝒩𝒫; specifically, it is unclear if checking if a degenerate circuit is minimal can always be done in polynomial number of steps in the number of neurons.

#### 2.5 Problem 3: Determine a Minimum Degenerate Circuit

We first formally state this question and it’s decision version.

##### Definition 14

(MinimumDC problem). *Determine a Minimum Degenerate Circuit*.

##### Definition 15

(MinimumDC (decision) problem). *For a given positive integer k, is there a Minimum Degenerate Circuit of size k?*

We now state the Cook reduction from the decision problem to the DC problem.

##### Proposition 3.

*MinimumDC (decision)* 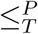 *MinimumDC*

As before, the reduction entails solving the MinimumDC problem, counting the number of neurons in the returned circuit, and returning a *Yes*, if the count is equal to *k* and *No* otherwise.

Next, we prove that the MinimumDC (decision) problem is 𝒩𝒫-hard by establishing a Karp reduction from CLIQUE. This implies that no general sub-exponential algorithms exist for any of the problems participating in the aforementioned reductions, unless 𝒫 = 𝒩𝒫.

##### Lemma 3.

*MinimumDC(decision) is 𝒩𝒫-hard*.

The reduction that was used to establish that DC(decision) is 𝒩𝒫-hard also turns out to imply that MinimumDC(decision) is 𝒩𝒫-hard. This is because, as proved in that construction, the (*k* + 2) sized degenerate circuit is also a minimum one, since the constructed circuit cannot, by design, have a (*k* + 1)-sized or smaller degenerate circuit. As a result, the constructed circuit has a (*k* + 2)-sized Minimum Degenerate Circuit, if and only if the graph has a *k*-clique.

Note that we are not asserting that MinimumDC(decision) is 𝒩𝒫-complete, since it is unclear if this problem is in the complexity class 𝒩𝒫; specifically, it is unclear if checking if a circuit is a minimum one can be done in a polynomial number of steps.

### 2.6 Problem 4: Determine the set of 1-Vital sets (or indicate if there are no 1-Vital sets)

It is straightforward to show that the set of 1-Vital Sets consists individually of neurons present in the intersection of all degenerate sets^16^. We first formally define the above problem.

#### Definition 16

(1-VITAL SETS). *Determine the set of 1-Vital sets*.

By definition, the problem above will return an empty set, if there are no 1-Vital Sets.

In contrast to previous problems, for technical reasons, here, we directly show a Cook reduction from CLIQUE to 1-VITAL SETS. This implies that no general sub-exponential algorithms exist for 1-VITAL SETS, unless 𝒫 = 𝒩𝒫. The proof below only describes the simplest construction. Afterward, we describe ways to further generalize the construction.

#### Lemma 4.

*CLIQUE* 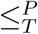 *1-VITAL SETS*

*Proof.* Given an undirected graph, for *k < n*, the reduction constructs a neural circuit (Figure S2) as described below, with the guarantee that the graph has a *k*-clique if and only if the set of 1-vital sets for the neural circuit is of size between 2 through *k* + 2; if the graph has no *k*-clique, then the set of 1-vital sets is of size *n* + 2, where *n* is the number of vertices in the undirected graph. If *k* = *n*, the reduction merely checks if the graph is a complete graph, and returns the answer. The Cook reduction is structured likewise.

The reduction is somewhat reminiscent of the reduction used to prove that DC(decision) is 𝒩𝒫-hard, although there are several differences. In the present case, if the undirected graph has no *k*-clique, then the circuit has exactly four degenerate circuits. These are the entire circuit, the circuit consisting of the interneuron circuit, plus the sensory and motor neurons, and the circuits obtained by the previous circuit plus each one of the coincidence-generator neurons. The intersection of these four DCs is the entire interneuron circuit plus the sensory and motor neuron, due to which the set of 1-vital sets is of size *n* + 2. On the other hand, if the undirected graph has exactly one *k*-clique, then the circuit has exactly five degenerate circuits. These are the aforementioned four DCs and, finally, the circuit formed by the neurons corresponding to the *k*-clique plus the sensory, motor and the coincidence generator neurons. As a result, the intersection becomes the *k* neurons in the interneuron circuit corresponding to the *k*-clique neurons plus the sensory and motor neurons – thus, the size of the set of 1-vital sets is *k* + 2. In case of undirected graphs with multiple *k*-cliques, in addition to the four degenerate circuits first described above, there is one degenerate circuit (corresponding to the fifth DC in the above account), for each such *k*-clique. As a result, it is straightforward to see that the intersection of the degenerate circuits comprises neurons that correspond to the vertices in the intersection of all the *k*-cliques in the graph, plus the sensory and motor neuron. Due to this, the set of 1-vital sets can be between size 2 through *k* + 2, if the undirected graph has one or more *k*-cliques.

**Figure S2:**
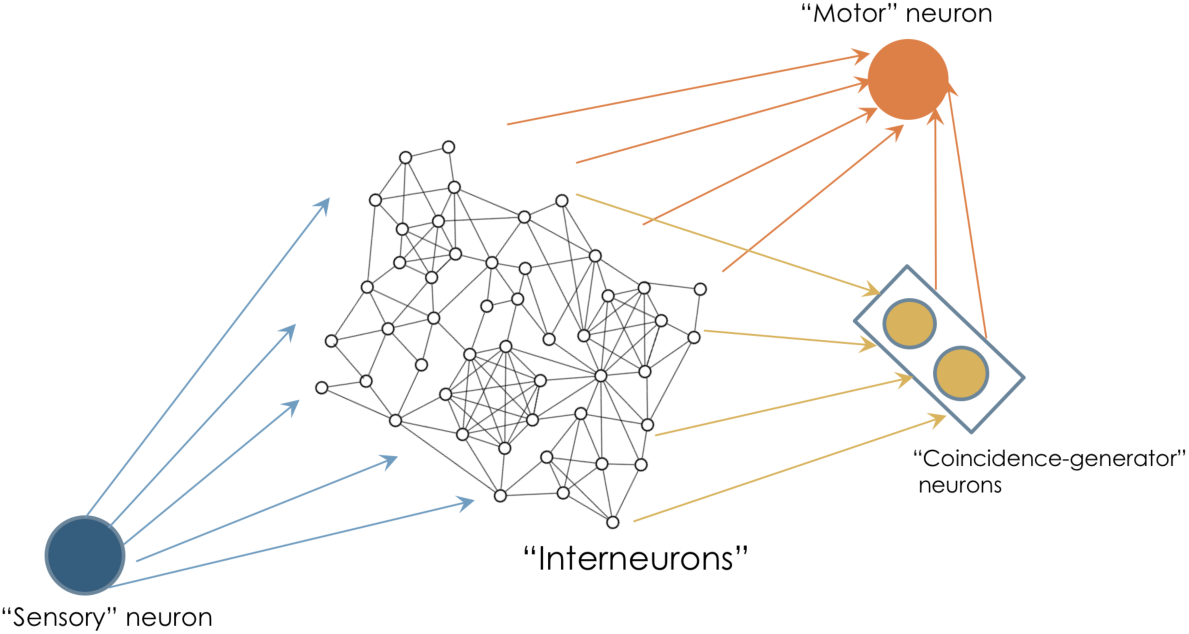
A schematic of the neural circuit that the reduction from CLIQUE to 1-VITAL SETS constructs, given any undirected graph. The interneuron circuit is what differs in the neural circuit for each given undirected graph.

We now describe the construction of the circuit and thereafter establish that it has the afore-mentioned properties. As before, the neural circuit constructed, has a single “sensory” and a single “motor” neuron. The sensory neuron signals the arrival of the pertinent stimulus by firing a single spike and the motor neuron, likewise, signals execution of the said behavior by firing a single spike (Figure S2). As before, the “interneuron” circuit connecting the sensory and motor neuron is constructed from the undirected graph by having an excitatory (inter) neuron in place of each graph vertex and bi-directional connections^17^ whenever there is an edge between vertices. Every interneuron receives input from the sensory neuron and is in turn connected to the motor neuron. In addition, all the interneurons are connected to a pair of “coincidence-generator” neurons which in turn are connected to the motor neuron. Conduction delays are the same within each pair of the following neuron types: sensory-inter, inter-inter coincidence-generator-to-motor and inter-motor. We consider here the state of the network, when all neurons start from quiescence. As before, the synapses connecting the sensory neuron and the interneuron are strong, so that a single sensory neuron spike causes an interneuron spike. Interneuron-to-motor neuron synapses are weak; concurrent spikes from all interneurons is sufficient to cause the motor neuron to spike. However, (& contrary to the previous construction), concurrent spiking from any (proper) subset of interneurons is insufficient to cause the motor neuron to spike, unless the coincidence-generator neurons concurrently send coincident spikes to the motor neuron; the coincidence-generator neurons do so, only in the case when they receive a pair of volleys from the interneurons, each having *k* spikes. The motor neuron is a coincidence detector, with respect to spikes received from these coincidence-generator neurons. As before, if an interneuron is not receiving input from the sensory neuron, it needs concurrent spiking from at least (*k* - 1) interneurons to produce a spike. We now describe the operation of the coincidence-generator neurons. This pair of neurons, on receiving a pair of volleys with *k* spikes apiece, will send coincident spikes^18^ to the motor neuron. For brevity, we will refer to them as the left and the right neuron respectively. The right neuron receives this input with a delay equal to the delay between the volleys plus some *c*, where *c* is the set up so that the two neurons fire coincident spikes for a pair of volleys with *k* spikes each. The postsynaptic potentials (PSPs) of both neurons have different rise slopes, so that other volley pairs do not result in coincident spikes output by them^19^. The coincident spikes will correspond to the second spike fired by the left neuron and the first spike fired by the right neuron. This is because the right neuron receives the first volley with a delay that allows the two neurons to, in a sense, “compare” the size of the two volleys.

The synaptic responses are set up in order to result in the following properties: (a) The entire circuit forms a degenerate circuit for the behavior. (b) The undirected graph has a *k*-clique, if and only if, for each *k*-clique, the circuit formed by the corresponding interneurons plus the coincidence-generator neurons, sensory and motor neuron form a degenerate circuit of size (*k* + 4) for the behavior.

Property (a) follows immediately from the construction. As for Property (b), it follows from the observation that a pair of volleys of *k* spikes apiece is fired if and only if (i) the degenerate circuit has exactly *k* interneurons as a subset and (ii) if those *k* interneurons form a clique. If the degenerate set has more (or less) than *k* interneurons^20^ the first volley will have more (or less) than *k* spikes. Secondly, if the *k* interneurons in the degenerate circuit don’t form a clique^21^, then the second volley will have less than *k* spikes. Property (b) follows, as a result.

We now briefly remark on ways to further generalize the construction. It turns out that the generalizations proposed for the DC (decision) reduction, with minor modifications, also work for the present reduction. The construction above has bi-directional connections between every pair of interneurons that is connected, which is thought to be not the case, physiologically. Likewise, all interneurons in the above construction are excitatory. Both these can be mitigated by the following addition to the above construction. Introduce arbitrary number of auxiliary interneurons (excitatory/inhibitory) with the constraint that each of them may receive connections or send out connections to at most (*k* - 2) other interneurons, which may be chosen arbitrarily. The inhibitory neurons do not send connections to the motor neuron or the coincidence-generator neurons. As a result, one has a neural circuit where the proportion of connections which are bidirectional in relation to unidirectional connections can be made rather small. Similarly, the circuit could also have proportion of excitatory and inhibitory neurons consistent with current estimates. The constraint described above on the number of connections ensures that the circuit has a (*k* + 4) sized degenerate circuit, if and only if the undirected graph has a *k*-clique and that the rest of the circuit properties established above are still correspondingly true.

#### 2.7 Problem 5: How many minimal ***k***-Vital Sets exist?

We first formally define the above problem.

##### Definition 17

(#MINIMAL-k-VITAL-SETS). *How many minimal k-Vital Sets exist?*

As with the previous problem, we directly show a Cook reduction from CLIQUE to #MINIMAL-k-VITAL-SETS. This implies that no general sub-exponential algorithms exist for #MINIMAL-k-VITAL-SETS, unless 𝒫 = 𝒩𝒫. The proof below only describes the simplest construction. Afterward, we describe ways to further generalize the construction.

##### Lemma 5.

*CLIQUE* 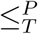 *#MINIMAL-k-VITAL-SETS*

*Proof.* Given an undirected graph, for *m < n*, the reduction constructs a neural circuit as described below, with the guarantee that the graph has an *m*-clique if and only if the number of minimal *k*-vital sets for the neural circuit is of size between 2*^k^* through *m^k^*; if the graph has no *k*-clique, then the number of minimal *k*-vital sets is of size *n^k^*, where *n* is the number of vertices in the undirected graph. If *m* = *n*, the reduction merely checks if the graph is a complete graph, and returns the answer. The Cook reduction is structured likewise.

The reduction is similar to that for 1-VITAL-SET, in that one has *k* modules in parallel, where each module is composed of the interneuron circuit plus the coincidence-generator neurons. Each module receives input from the sensory neuron with a different delay; this is in order to ensure that activity from each module does not “interfere” with activity from any other module insofar as concurrently influencing the output of the motor neuron. Furthermore, there is inhibition from the motor neuron to each module, that ensures that the motor neuron only fires once for each spike from the sensory neuron.

Choosing a 1-Vital Set neuron (as implied by the construction of Problem 4) from each module and doing so for all the *k* modules gives us a *k*-vital set for the present circuit. This is because, on doing so, none of the modules can individually cause the motor neurons to spike and by construction, they cannot co-operatively do so either. Furthermore, each such *k*-vital set is minimal because removing any neuron from it will cause the corresponding module to be able to cause the motor neuron to spike, on receiving a sensory neuron spike. Now, from Problem 4, if the graph has a *m*-clique, each module will identically have between 2 and *m* 1-vital sets in the sense of Problem 4. As a result, combinatorially, the number of *k*-vital sets for the present problem is obtained by independently choosing a “1-vital set” from each module; there are between 2*^k^* through *m^k^*such *k*-vital sets. Likewise, if the graph does not have an *m*-clique, the constructed neural circuit has number of *k*-vital sets equal to *n^k^*. Finally, no other *k*-vital sets exist, in addition to those enumerated above. This arises out of two observations. Firstly, every minimal *k*-vital set needs to have exactly one neuron from each module; if not there would exist a module with all neurons intact, which would cause it to form a degenerate circuit. Secondly, the neuron so chosen from each module has to be 1-vital set in the sense of Problem 4, otherwise there would exist a way to form a degenerate circuit in the sense of Problem 4 and as a result for the present problem as well. This completes the proof.

The analogous generalizations suggested for Problem 4 work here as well.

An important observation in the current reduction, is that the constructed circuit is highly modular. In fact, with the suggested generalizations, it can have each module looking different from the others to an arbitrary degree. It has been previously hypothesized (*27*) that modularity could alleviate the combinatorial complexity of understanding biological systems. This result represents an important demonstration of a question about a modular circuit that cannot be answered with a tractable number of experiments as well, unless 𝒫 = 𝒩𝒫.

##### On large cliques

One observation is that the neural circuits constructed in many of the aforementioned reductions can have fairly large cliques. This is not thought to be the case in nervous systems; for example, circuit reconstructions in the Blue Brain Project have fairly small-sized cliques (*31*) and this is indeed the case with several random graph models as well (*32*). We now outline a modification to the construction which results in the elimination of large cliques in the neural circuits constructed, while retaining the stated property with respect to the degenerate circuits in the aforementioned problems.

Briefly, the idea is to substitute edges in the neural circuits constructed previously, with small neural circuits which, in effect relay an incoming spike to an outgoing spike. Further, they have the property that silencing any neuron within the subcircuit disrupts this relay process. None of the neurons in this subcircuit send their output to the motor neuron. Each such subcircuit has the same number of neurons, *m*, although they could each be a different circuit with the properties stated above. As a result, instead of the presence of a degenerate circuit of size *k* + 2, we have a degenerate circuit of size 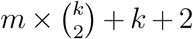, whenever the undirected graph has a clique of size *k*. For technical reasons, in the case of Problems 1 through 3, however, we will have to start from the neural circuit constructed in Problem 4 (which also works on its own, for Problems 1 through 3). It is straightforward to see that this modification guarantees that if the undirected graph has a clique of size *k*, then the neural circuit constructed with this modification has a degenerate circuit of size 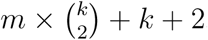, without the presence of cliques, unless they are present in the aforementioned subcircuits. We, however, also need to ensure that the converse is true. The only case, where the converse may not be true is if, for appropriately large *m*, 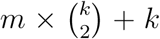 is greater than or equal to *n*, where *n* is the number of vertices in the undirected graph. In this case, one might have a degenerate circuit made up only of the corresponding *n* interneurons that send input to the motor neuron, plus the sensory and motor neuron. To eliminate this possibility, the final step of the modification is to add 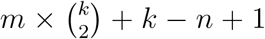 auxiliary interneurons which receive input from the sensory neuron and in turn send input to the motor neuron, with the stipulation that the motor neuron spikes (given a single volley from the interneurons) only if these auxiliary neurons plus the aforementioned *n* interneurons spike concurrently. These neurons may receive input from other interneurons or send their outputs, with the constraint that each neuron sends or receives spikes from no more than *k* - 2 other interneurons. This ensures that the converse is also true.

In closing, the above modification establishes that the presence of modest size cliques in the constructed neural circuits isn’t a necessary feature for the reductions to be true.

#### 2.8 Problem 6: Determine a minimal ***k***-Vital Set, or indicate if none exists of that size

We first formally state the question and it’s decision version.

##### Definition 18

(MINIMAL VITAL SET). *For a given positive integer k, determine a minimal k-Vital Set, or indicate if none exists of that size*.

##### Definition 19

(MINIMAL VITAL SET (decision)). *For a given positive integer k, does there exist a minimal k-Vital Set?*

We now state the Cook reduction from MINIMAL VITAL SET (decision) to MINIMAL VITAL SET.

##### Proposition 4.

*MINIMAL VITAL SET (decision)* 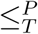 *MINIMAL VITAL SET*

The reduction simply entails solving the MINIMAL VITAL SET problem for the same inputs and examining if the solution returned is a degenerate circuit of size *k* or if it is indicated that there is no such degenerate circuit; in the former case the decision version returns a *Yes* and in the latter case it returns a *No*.

In order to show that MINIMAL VITAL SET (decision), and consequently MINIMAL VITAL SET, have no sub-exponential algorithms, unless 𝒫 = 𝒩𝒫, we start from the HITTING SET problem, which is another known 𝒩𝒫-complete problem, reduce it to another problem that we will define, which will then be reduced to MINIMAL VITAL SET (decision).

We first state the HITTING SET problem, which was one of the first to be shown (*33*) to be 𝒩𝒫-complete. Given a finite set *U* and a collection *C* of its subsets, a *hitting set* is a set of elements from *U*, such that it “hits”, i.e. has a non-null intersection with, every set present in *C*. A hitting set of size *k* is called a *k*-Hitting Set. A hitting set, none of whose proper subsets are also hitting sets is called a *minimal hitting set*.

##### Definition 20

(HITTING SET). *Given a finite set U, a collection C of its subsets and a positive integer k, does there exist a subset H of U, with* |*H*| ≤ *k, such that H has within it, at least one element from each subset present in C*.

We now define another problem, namely one of determining if there exists a minimal hitting set of a certain size.

##### Definition 21

(MINIMAL HITTING SET). *Given a finite set U, a collection C of its subsets and a positive integer k, does there exist a subset H of U, with* |*H*| = *k, with the property that H has within it, at least one element from each subset present in C, and furthermore that no proper subset of H has this property*.

The two problems above are related in that there exists a Cook reduction from the former to the latter, which we now state and describe.

##### Proposition 5.

*HITTING SET* 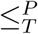 *MINIMAL HITTING SET*

The reduction entails solving the MINIMAL HITTING SET problem, while running the parameter *k* of MINIMAL HITTING SET from 1 through *k*; the HITTING SET problem has a *Yes* answer if and only if at least one those problem instances of MINIMAL HITTING SET has a *Yes* answer. This Cook reduction implies that there exists no general sub-exponential algorithm for MINIMAL HITTING SET, unless 𝒫 = 𝒩𝒫.

Next, we establish a Karp reduction from MINIMAL HITTING SET to MINIMAL VITAL SET (decision).

##### Lemma 6.

*MINIMAL HITTING SET* 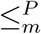 *MINIMAL VITAL SET (decision)*

*Proof.* We first describe the construction of a neural circuit, given a finite set *U*, a collection *C* of its subsets and a positive integer *k >* 1; if *k* = 1, the reduction simply checks and answers if the MINIMAL HITTING SET instance has a minimal hitting set of size 1. The construction will guarantee there there exists a minimal *k*-Hitting Set for the aforementioned instance, if and only if the neural circuit has a minimal *k*-Vital Set.

The neural circuit has a sensory and motor neuron, similar to previous constructions in that the sensory neuron fires a single action potential to signal the arrival of the stimulus and that the motor neuron fires a single action potential to signal the execution of behavior. The sensory neuron sends information to the motor neuron via two successive layers of neurons. The first layer (i.e. the one closer to the sensory neuron) has one neuron corresponding to each element in *U*. Every *U* neuron receives input from the sensory neuron via a strong synapse such that a single sensory neuron spike elicits a spike in that neuron. The second layer has (*k* + 1) neurons corresponding to each subset in *C* of the MINIMAL HITTING SET instance. Each such neuron gets input from the neurons of layer 1 that correspond to elements from *U* in its subset. Each Layer 2 neuron fires if and only if all Layer 1 neurons it is connected to fire concurrently. Finally, each layer 2 neuron connects to the motor neuron, such that firing of any one (or more) Layer 2 neuron is sufficient to cause the motor neuron to spike.

Due to the construction, the neural circuit has a minimal *k*-Vital Set, if and only if there is a minimal *k*-Hitting set in the MINIMAL HITTING SET instance, as we now prove. One direction is showing that if there is a minimal *k*-Hitting Set, then there is a minimal *k*-Vital set, which is straightforward from the construction. The minimal *k*-Vital Set consists exactly of the neurons in the first layer that correspond to elements that form a minimal *k*-Hitting Set. This is because silencing those neurons leads to every neuron in Layer 2 not receiving input from at least one neuron in Layer 1 that it is connected to, resulting in it not producing a spike, upon the sensory neuron first firing a spike. As a result, the motor neuron cannot fire a spike, either. Secondly, it is a minimal Vital Set, since any subset of it, by definition would not correspondingly form a Hitting Set, due to which there is a subset of *C* that doesn’t intersect with it. This results in at least (*k* + 1) neurons in Layer 2 that end up receiving input from all Layer 1 neurons that they receive input from, which results in the said set not being a Vital Set. Next, we need to show that if there is no minimal *k*-Hitting Set, then there is no minimal *k*-Vital Set. If there did exist one, such a *k*-Vital Set cannot involve the sensory or motor neuron, since any minimal Vital Set involving them is a 1-Vital Set. It cannot just involve *k* first-layer neurons, since that would imply that the corresponding elements from the set constitute a minimal *k*-Hitting Set, which isn’t the case. Finally, if a second layer neuron is an element of a minimal *k*-Vital Set, then every such neuron, which is receiving input from the same first-layer neurons must also be in the minimal *k*-Vital Set for reasons of symmetry. Since there are (*k* +1) such neurons in total, corresponding to each such subset in *C*, the minimal Vital Set is of size at least (*k* + 1), which is not equal to *k*. Together, this implies that there is no minimal *k*-Vital Set, if there is no minimal *k*-Hitting Set. This completes the proof.

The above Karp reduction implies that there exists no general sub-exponential algorithm for MINIMAL VITAL SET (decision), unless 𝒫 = 𝒩𝒫. As a result, there exists no general sub-exponential algorithm for MINIMAL VITAL SET, unless 𝒫 = 𝒩𝒫, since we showed a Cook reduction from MINIMAL VITAL SET (decision) to MINIMAL VITAL SET.

### 3 Quasi-minimal degenerate circuits and a Θ(log *n*) algorithm to determine one

The idea of isolating subsets of neurons which causally participate in computations leading to the behavior is a potentially useful one. Having found such subsets, one could then go in and find out detailed mechanisms by which the said subset of neurons perform the computations that lead to behavior. However, what we have demonstrated is that for many such notions of participation, no general algorithms exist that use sub-exponential number of experiments, unless 𝒫 = 𝒩𝒫. Furthermore, as discussed in the main text, the feasible regime of experiments is one where number of experiments that one can perform scales sub-linearly in the number of neurons. Therefore, for many such circuits, it might be infeasible to determine the aforementioned types of subsets.

One approach is to somewhat relax the notions of participation to define types of subsets that can always be determined with sub-linear experiments in the number of neurons, no matter what class of neural circuits is involved. Such subsets still contain neurons that causally participate in computation, although they might have neurons that are not involved as well^22^.

In that spirit, we present here a notion of a quasi-minimal degenerate circuit that, conceptually, is stricter than just a (vanilla) degenerate circuit, but isn’t quite a minimal degenerate circuit. We then present an algorithm to determine a QMDC that requires a logarithmic number of experiments in the number of neurons in the circuit. A logarithmic function scales extremely slowly with input size. The logarithm^23^ of a 100 billion is about 37. That is, to determine such a set for a brain with a 100 billion neurons, one would need about 37 experiments – an eminently tractable number. Binary search or its variants are arguably the most used logarithmic-time algorithms in the digital world today and are largely responsible for seemingly instantaneous search on large databases that characterize a lot of activity on the internet today.

Before going ahead and defining the notion of a Quasi-minimal Degenerate Circuit, we first motivate it. In general, for a behavior with the readout being correctly elicited in an intact brain, if one wants to find a degenerate circuit, the entire brain would be it and therefore asking for any degenerate circuit has a trivial answer. On the other hand, asking for a degenerate circuit of a certain size, in general, requires exponentially-many experiments in the number of neurons, unless 𝒫 = 𝒩𝒫 (cf. Problem 1). We would like to have a notion that is more constrained than just being a degenerate circuit, yet, so that it can be determined with sub-linear number of experiments. This, broadly speaking, is the motivation in defining the notion of a Quasi-minimal degenerate circuit.

Consider a nested sequence of subsets of neurons starting from all the neurons to the empty set, with the next set in the sequence, constructed from the previous set by removing one neuron^24^. As one goes from the beginning to the end in the sequence, associate a label 1 with the current set if it is a degenerate circuit^25^ and a label 0, if it is not. One might imagine that in moving through the sequence, we will initially see several 1-sets, possibly interspersed with 0-sets. Later in the sequence, one expects to see mostly 0-sets. We seek to find a 1-set with a 0-set immediately on its right. This would correspond it to being minimal in a certain sense^26^ with respect to the neuron that is silenced to get to the 0-set. We call such a 1-set a *Quasi-minimal Degenerate Circuit (QMDC)*. Observe that there is at least one Quasi-minimal degenerate circuit, if the entire brain is a degenerate circuit and if the empty set (i.e. with all neurons silenced) is not a degenerate circuit.

The algorithm to determine a QMDC is an adaptation of standard binary search. There are two pointers, which we will call the 1-pointer and the 0-pointer. The 1-pointer is initially positioned at the first set in the sequence and the 0-pointer is positioned at the last set in the sequence, which is the empty set. At each step one considers the set which is midway in the sequence between the current 1-pointer and the current 0-pointer and runs an experiment to check if it is a 1-set or a 0-set (i.e. if it is a degenerate circuit or not). If it is a 1-set, the 1-pointer is re-positioned at that set, otherwise the 0-pointer is re-positioned at that set. One iterates through this process until the 0-pointer ends up being immediately after the 1-pointer. At this stage, the set corresponding to the 1-pointer is a QMDC and the algorithm halts.

The analysis for how many steps this algorithm takes is largely identical to that of standard binary search. The key observation is that the size of the sequence processed by the algorithm halves at each iteration as a result of which it ends up using approximately log_2_ *n* number of experiments, where *n* is the total number of neurons.

## 4 Conclusion

Empirically understanding how neural circuits mechanistically cause behavior, seems to be a prerequisite, for distilling the principles that govern their operation, or to validate said principles post hoc. The current technological renaissance at the circuit level in experimental neuroscience is largely aimed at affording us the means to obtain such an empirical understanding. In such experimental settings, multiple experiments are usually necessary to arrive at a causal understanding of circuit mechanisms. It has been unclear *how many* experiments are needed to arrive at a comprehensive empirical understanding of how neural circuits mechanistically cause behavior.

The results here suggest the existence of an algorithmic barrier to obtaining such an understanding. Specifically, there exist questions about mechanistic circuit computation, for which no general circuit interrogation algorithms exist that are guaranteed to answer said questions with a tractable number of experiments, in general. A related, if subtle, implication is that notions of understanding that might allow us to answer such questions in short order themselves cannot, in general, be tractably acquired. While we have demonstrated *six* such questions here, there likely exist many more such questions that cannot be tractably answered, in general. Secondly, we have demonstrated that the feasible regime of experiments, for most organisms of interest, is one in which the number of experiments one can perform, scales sub-linearly in the number of neurons. This remarkable gap between the worst-case and the feasible suggests strong constraints on notions of understanding that might be experimentally establishable in practice.

It is also worth recalling that the analysis here has treated the case of a single behavioral readout, for a single state. In practice, we will need to understand neural circuits with respect to multiple behavioral readouts and during multiple states, which further increases the number of experiments necessary to so understand the system. Also, we have set up our notions of neurons “participating” in mechanistic computation without considering temporal aspects of participation. That is, sets of neurons might be crucial for mechanistic circuit computation at specific times in course of expressing the behavior, and not others. Establishing this temporal aspect will likely require more experiments as well. Furthermore, the analysis here has focused on the question of how neural circuits *acutely* cause behavior. Recent beautiful work (*34*) has demonstrated instances of neural circuits that are acutely necessary for behavior, yet are not so required chronically. As a result, a distinction has been proposed between permissive and instructive circuits that cause behavior. Empirically establishing such distinctions seems to add an additional layer of complexity that will likely require more experiments as well.

Now, one might make the argument that the approach taken here of using whole-brain circuit interrogation to comprehensively understand the neural mechanisms of a behavior is excessive; that is, it should suffice to just study how the relevant brain region(s) cause the behavior instead. While, classically, nervous systems have been parcellated into regions based on putative function, this is a view that is increasingly being challenged in many cases. Specifically, regions thought to be important for certain types of tasks have been shown to be dispensable for said tasks and vice versa. For example, it has recently been shown (*35*) that the motor cortex is not necessary for the execution of certain learned movements and that the barrel cortex is not necessary, both acutely and chronically, for certain behaviors involving whisking and somatosensation (*36*). In contrast, the cerebellum has recently been shown to be indispensable for maintaining short-term memory in a delayed sensory discrimination task (*37*) as well as in an evidence-accumulation-based decision-making task (*38*), in addition to the pre-frontal cortex (PFC). Whereas, previous accounts of the neural substrate of short-term memory (in such cases) have largely focused on the PFC, going forward, any comprehensive account ought to also detail the role of the cerebellum. In short, our understanding of such putative function is ongoing and might involve several subtle task-dependent distinctions that involve neurons in brain regions that have been heretofore unexplored for said behaviors. Furthermore, the functional roles of many areas aren’t even particularly clear. For example, the functional role of the claustrum has been largely unclear (*39*) (see also, for example some recent early work in this direction (*40–42*)), in spite of the fact that it is the most densely connected structure by volume in the human brain (*43*) and has been hypothesized famously by Crick and Koch (*44*) to be the seat of consciousness. For these reasons, studying and interrogating whole-brain activity, where possible, is a reasonable way of arriving at a comprehensive understanding of circuit mechanisms and we are increasingly approaching the means to do so (*45–48*) in several organisms. Indeed, this paradigm of concurrently imaging and perturbing activity while monitoring behavior has already been fruitfully employed in larval zebrafish (*49, 50*) to study causal effects of neurons on behavior. Moreover, even if we were somehow able to a priori isolate the causal substrate of the behavior to being present in a small subset of the brain (e.g. 1%), it would still be prohibitive to run exponentially-many experiments in that subset, for most organisms of interest. Therefore, the issue of ascertaining tractable algorithms for understanding how neural circuits mechanistically cause behavior remains a pertinent question, even in this case.

Another argument that might be made is that the “right” level for understanding of neural circuit mechanisms is with respect to cell types, as opposed to individual cells. Indeed, many current circuit interrogation tools allow us to easily manipulate cells of certain genetically-identified cell types. However, multiple types of circuit computations and behavior (*51–54*) have been shown to depend on neurons that are genetically-similar, but spatially intermixed. In fact, this has motivated the development of techniques such as two-photon optogenetics (*55*) and holographic patterning techniques (*56*) that offer the capability of perturbing arbitrary subsets of neurons.

The presence of modularity in nervous systems has been hypothesized to potentially help tame the number of experiments needed to analyze them (*27*) and indeed there appears to be emerging evidence of modularity in nervous systems as exemplified by recent connectomic data (*57*). While this hypothesis is compelling, this does bring up some issues. The first is the question of identifying such modules, especially if they are not identical but degenerate (*58*). Secondly, the modules themselves then need to be understood, and this might be onerous especially if they are large. Finally, the presence of modules, in and of itself, does not make intractable algorithmic problems go away, as the reduction for Problem 5 demonstrates. In that reduction, we construct a modular circuit, initially with identical modules, yet for which Problem 5 cannot be answered with sub-exponential number of experiments in the number of neurons, unless 𝒫 = 𝒩𝒫. After the simplest construction of the neural circuit, we suggest modifications, as a result of which the modules can be different from each other to an arbitrary extent. The algorithmic intractability result continues to exist in these cases. We therefore suggest that while the modular case may make the problem more tractable to solve in many cases, that conclusion is not foregone and further work is needed to study this case more carefully.

Classically, the prevalent conceptual framework for understanding neural circuits has been the Marr levels framework (*59, 60*). Briefly, Marr proposed that the nervous system needs to be understood at three levels^27^ – computational, algorithmic and implementational. The computational level pertains to the problems that the system is trying to solve, the algorithmic level is about the algorithms used to solve the problems and the implementational level deals with how the algorithms are implemented in the neural substrate. Indeed, this compelling framework has shaped the thinking in the field for over four decades. However, more recently, questions have emerged about the utility of this framework, especially in relation to its bottom two levels, articulated, notably by Stephane Mallat (*61*). The crux of the issue is that the levels framework implicitly assumes that there exists a clean separation between the algorithmic and implementation levels, which may not be the case. For example, Computer Vision has looked to the brain as a proof of principle and a great deal of the early work was inspired by an understanding of the mammalian visual system at the time. In particular, Computer Vision has been based on the premise that succinct vision algorithms in nervous systems (and elsewhere) exist independent of their neural substrates and therefore it ought to be possible to determine such algorithms, even if they are not inferred from nervous systems. While classical Computer Vision has enjoyed some successes, it is safe to say that most of the fundamental problems were not satisfactorily solved in practice, in spite of decades of intense work that was grounded on this premise. The recent astounding success of Deep Learning in Computer Vision, therefore, has challenged this premise. In fact, while contemporary deep learning systems perform better than humans in tasks such as visual recognition, we have failed to distill succinct algorithms from these networks (that exist independent of the neural substrate). Indeed, we presently don’t even have an adequate understanding of the working of these deep networks, in spite of our ability to manipulate them at will. It is possible that an analogous algorithmic barrier exists in comprehensively understanding deep networks as well. Getting back to neural circuits, it thus seems possible that the algorithmic level of understanding cannot always be cleanly separated from the implementational level, which likely exacerbates the problem of understanding neural circuits succinctly.

As a consequence of this work, we argue that a sustained effort is necessary to investigate the issue of tractability of neural circuit interrogation algorithms for causal understanding of neural circuits. Such an effort could envision building a suite of questions about neural circuit computation, with the questions being enumerated in increasing order of number of experiments required. This is a resource that experimentalists could directly tap into, in designing their experiments. As discussed in the main text, in general, there are two directions to take here. The first, is one of determining the questions^28^ about neural circuit understanding that, in fact, always need only sub-linear number of experiments to answer. As a first example, we have provided one such question and a corresponding algorithm here – namely for determining a Quasi-minimal Degenerate Circuit. The second direction is to design algorithms that are guaranteed to run with a small number of experiments for certain sub-classes of neural circuits. Theory has an important role to play here in that it is often an implicit way to specify such sub-classes of neural circuits. For example, if we know that a certain behavior is mediated by synfire chains, one can immediately infer the presence of a number of vital sets – which would correspond to sets of neurons, which when silenced, cause the propagation of synchronous activity to be abolished. A larger related implication that the results here suggest, is that theory-agnostic neural circuit interrogation might end up being especially onerous, if not outright infeasible, in many cases.

Finally, we could not help but observe that the problem of understanding computation in brains is analogous to a similar question in Computability Theory that goes back to the work of Turing and others. Briefly, Turing Machines have finite descriptions, due to which it is, in principle, possible for one Turing Machine to take as input a description of another Turing Machine and seek to answer questions about computation in the latter Turing Machine. Alan Turing (*62*) was the first to show an example of a question in this setting that is undecidable; that problem has since been referred to as the Halting Problem. In subsequent landmark work, Rice (*63*) with his eponymous theorem proved that answering a large (and interesting) class of such questions for the (former) Turing Machine is undecidable. For Computer Science, this implies, for example, that one cannot have automated debugging programs (or algorithms) that are guaranteed to always work correctly. Undecidability in the bounded case often manifests as 𝒩𝒫-hardness (*33*), which parallels the results established here. In short, there likely exist deeper connections from the mathematical foundations of Computer Science, to the question of understanding mechanistic computation in neural circuits, which merit further investigation.

## Footnotes

1 Imaging of neural circuit activity alone, will only tell us which neurons’ activity is correlated with aspects of stimulus/behavior. In order to determine causal substrates, it is necessary, in general, to perform (multiple) perturbational experiments.

2 We emphasize that doing so, is *not* a simplifying assumption. Indeed, algorithmic constraints apply not just to computers but also to humans that design experiments.

3 See Supplementary Text for more on this point.

4 If, remarkably, 𝒫 = 𝒩𝒫 were true, it would mean that hundreds of computational problems – many of them commercially important and extensively studied for decades – would have sub-exponential algorithms, where none have been found to date.

5 The hypothesis mentioned in the abstract (that is widely thought to be true) is that 𝒫 ≠ 𝒩𝒫.

6 One chooses to study and understand circuits specific to a single individual, since there is thought to be inter-individual variability across individuals of the same species.

7 This corresponds to obtaining an understanding for a single “state” of the system. Doing so, imposes a lower bound on understanding all states. Going forward, we will assume that the state is so fixed in each instance.

8 It is important to point out that notions of understanding described, are with respect to this readout. That is, the readout may choose to ignore many aspects of the behavior, which will not be subjects of such an understanding. Conceivably, one can construct other readouts that might so consider such aspects of behavior.

9 This is because there could exist another degenerate circuit identical to the present one, except for the absence of the neuron in question.

10 The questions are deliberately formulated, so that the answers cannot be exponentially-long in the number of neurons, for otherwise, just enumerating it would be exponential. e.g. asking to list all degenerate circuits could be exponential in the number of neurons.

11 Informally, a reduction is a recipe to quickly convert any instance of one problem (Problem A) into an instance of another problem (Problem B), such that a solution to Problem B can be quickly mapped back to a solution of Problem A as well. Therefore, the existence of an efficient algorithm for Problem B immediately implies the existence of an efficient algorithm for Problem A, via the reduction. More profoundly, if there exists no efficient algorithm for Problem A, the reduction implies that Problem B cannot have an efficient algorithm either; otherwise there would be a logical contradiction.

12 This is an outcome of the curious observation that most organisms are estimated to have more neurons in their nervous systems than the average number of seconds they live.

1 corresponding to different variants of Turing machines

2 The connectome can currently only be determined post-hoc. The results here will imply that even if the connectome were known in advance, there exist questions that need exponentially-manu experiments, in general, unless 𝒫 = 𝒩𝒫. The results, of course, also hold if the connectome is not known.

3 unless 𝒫 = 𝒩𝒫.

4 unless 𝒫 = 𝒩𝒫.

5 This is because *n* times a function that is sub-exponential in *n*, remains sub-exponential in *n*.

6 The vital set turns out to be closely related to the notion of a Hitting Set in Computer Science (& Discrete Mathematics). Specifically, the vital set as defined here is the hitting set of the set of all degenerate sets.

7 especially for the reader unfamiliar with Theoretical Computer Science.

8 named in honor of Stephen A. Cook. This type of reduction is also often called a polynomial-time Turing reduction.

9 named in honor of Richard M. Karp. This type of reduction is also often called a many-one polynomial-time reduction.

10 since search problems are not decision problems.

11 measured over a certain window.

12 Without this latter step, this problem would simply be shown to be 𝒩𝒫-hard.

13 As we will elaborate later, the appearance of a large number of bi-directional connections is not necessary for the construction to work. Indeed, the proportion of bi-directional connections to unidirectional connections can be made arbitrarily small, with the reduction still working.

14 Specifically, a pair of volleys here is defined as two sets of concurrent spikes which are separated temporally by time equal to the inter-inter conduction delay.

15 If not, then the *k* non-silenced interneurons have a clique, which contradicts the hypothesis.

16 This intersection is also identical to the intersection of all the minimal degenerate sets.

17 As we will elaborate later, the appearance of a large number of bi-directional connections is not necessary for the construction to work.

18 Here, coincident might be defined as being with some ϵ ms of each other, where ϵ is small.

19 within the aforementioned ɛ.

20 That is, if (i) is not true.

21 violating (ii).

22 It is important to remember that this is only a heuristic method. Indeed, this is not a route to always get to the aforementioned type of causal subsets, for otherwise, we would have 𝒫 = 𝒩𝒫.

23 base 2, in Computer Science, unless otherwise stated.

24 The one neuron that is removed may be arbitrarily chosen. One could also use heuristics such as choose the neuron with the highest firing rate among the intact neurons and so on. The algorithm is invariant to this choice.

25 As before, the set is a degenerate circuit, with respect to a fixed behavior (& associated binary behavioural readout) and state. One either needs to have the ability to have the circuit in the same state while performing each experiment or know that the set of degenerate sets is invariant to a certain class of states. The latter possibility could be operationally checked by performing multiple repeats of the experiment which will increase the number of experiments by a constant factor.

26 This is a less strict sense of minimality, since a set further ahead in the sequence could be a 1-set.

27 The first articulation of the levels framework (*59*) mentioned four levels, although the subsequent book and indeed the popular memory is of three levels.

28 It also remains to be investigated if there exist efficient approximation algorithms – algorithms that solve approximate versions of the stated questions – for the six questions stated. It would be quite remarkable, if there existed such approximation algorithms that only needed sub-linear experiments in the number of neurons. In general, even for other questions, it is conceivable that there exist efficient approximation algorithms even if there might not exist efficient algorithms for the exact version of the question.

